# Morphological and genetic confirmation of the millipede *Chordeuma sylvestre* C. L. Koch, 1847 new to Ireland

**DOI:** 10.1101/2020.05.17.100263

**Authors:** Willson Gaul, Andrew J Tighe

## Abstract

We report the millipede *Chordeuma sylvestre* C. L. Koch, 1847 for the first time from Ireland. We used morphology of male gonopods and DNA barcoding of the COI gene from two individuals to confirm the identification of one male and one female collected from the campus of University College Dublin. DNA barcoding revealed two mitochondrial haplotypes, implying that colonization was from multiple females or eggs from multiple females. To evaluate how likely it is that *C. sylvestre* arrived relatively recently in Ireland, we used species richness estimates and species accumulation curves to evaluate the completeness of the species list compiled during two previous periods of intensive millipede recording going back to 1971. Millipede recording in Ireland was far from complete during the period 1971 – 1984, with multiple species present but undetected. Recording was more complete during the period 1986 – 2005. The area within 20 km of the new *C. sylvestre* records was poorly recorded during the period 1986 – 2005, so we cannot rule out the possibility that *C. sylvestre* has been in Dublin for at least a few decades. However, the disjunct location of the current records compared to the species’s known native range in northern France, the urban location of the current site, and the lack of records from the relatively intensive millipede recording in Ireland between 1986 and 2005 suggest that *C. sylvestre* is a recent anthropogenic introduction in Ireland.

## Introduction

The millipede *Chordeuma sylvestre* C. L. Koch, 1847 is native to central Europe, and is known from a few locations in Great Britain (Gregory, 2016). It has not been previously recorded from Ireland. Kime (2004) did not consider *C. sylvestre* a habitat specialist and noted that it has been found in coniferous and deciduous woodlands, peat bogs, and moors, among other habitats. It therefore seems unlikely that *C. sylvestre* has been absent from Ireland because of environmental factors. Morphological identification of *C. sylvestre* requires examining adult males – females of *C. sylvestre* cannot be distinguished from females of the congener *C. proximum* Ribaut, which occurs in Ireland and Great Britain (Blower, 1985).

Here, we present records of *C. sylvestre* from Dublin, Ireland, including DNA barcoding identification of a female specimen for which morphological identification was not possible. To gain insight into whether *C. sylvestre* is a relatively recent introduction to Ireland or whether it has been present but undetected for many years, we examined Irish millipede records from 1971 to 2005. We evaluated the completeness of millipede species lists compiled during previous recording by using species accumulation curves and comparing estimated and observed species richness.

## Methods

### Specimen collection, preservation, and morphological examination

On 18 January 2020, the first author (WG) collected multiple millipedes from the campus of University College Dublin. Searching was done by turning and sifting leaf litter and soil with a small trowel and by hand, and collecting all millipedes encountered. Collected specimens were placed together into a vial with 70% isopropyl alcohol and stored at room temperature. On 6 March 2020 WG returned to the same location with the explicit purpose of looking for another *Chordeuma* specimen, and collected a female, which was placed by itself into a plastic vial and frozen (without being put in alcohol). The female specimen was kept frozen for three days until DNA extraction. All collected specimens were examined with a microscope using magnifications of 25 to 200 times. Male gonopods were dissected, slide mounted in Euparol, and photographed. Species identification was done by WG using Blower (1985) and the species pages on the British Myriapod and Isopod group website (BMIG, 2020, www.bmig.org.uk/home). Records with photographs were submitted to iRecord (www.brc.ac.uk/irecord/).

### DNA barcoding

On 9 March 2020, tissue samples of about 4 or 5 segments, including legs, exoskeleton, and internal organs, were taken from both the male specimen that had been stored in 70% isopropyl for two months and from the female specimen that had been frozen for three days. The tissue samples were separately placed in 400 µl of 10% Chelex solution in a 1.5 mL tube (Eppendorf) and crushed using a pestle (Eppendorf), followed by the addition of 12 µl of proteinase K (20 mg/mL) to each tube. The tubes were briefly vortexed and then incubated at 56°C for 2 hours. After incubation the samples were heated to 99°C for 15 minutes, followed by centrifugation for 1 minute at 20,817 G, and 50 µl of DNA supernatant was removed from the sample and transferred to a new tube.

In order to amplify the *cytochrome oxidase I* (COI) gene, the primers LEPF1 [5-ATT CAA CCA ATC ATA AAG ATA T-3] and LEPR1 [5-TAA ACT TCT GGA TGT CCA AAA A-3] (Hebert *et al.*, 2004) were used. A 25 μl PCR master mix was prepared in a UV-sterilized hood and consisted of 3.125 μl Buffer (Kapa Biosystems), 1.25 μl dNTP (Invitrogen), 1.25 μl of each primer (10 μM) (Integrated DNA Technologies), 0.125 μl Taq polymerase (Kapa Biosystems), 17 μl water, and 1 μl of DNA extract. PCR conditions were as follows: initiation at 94°C for 1 min, followed by 5 cycles of 94 °C for 40 s, 45°C for 40 s and then 72°C for 1 min, then 35 cycles of 94°C for 40 s, 51°C for 40 s and 72°C for 1 min. The final elongation step was 72°C for 5 min, after which samples were held at 4°C. All PCR amplifications were carried out in a separate room from where DNA was extracted. Successful amplification was based on the presence of a band of the correct molecular weight in a 1% agarose gel after electrophoresis. Both samples amplified successfully and were sent for commercial Sanger sequencing (Macrogen).

Clean, unambiguous sequence was received for both samples, and the forward and reverse strands were aligned using Geneious version 10.2.3 (Kearse *et al*., 2012). Strand alignment and trimming resulted in a 674 bp consensus sequence for the male specimen and 662 bp for the female. The two sequences were then aligned using ClustalX version 2.1 (Larkin *et al*., 2007). Both sequences were then separately analyzed using BLAST (https://blast.ncbi.nlm.nih.gov/Blast.cgi) to find matching sequences in GenBank.

### Completeness of millipede recording in Ireland

We assessed the completeness of millipede recording on the island of Ireland (Ireland and Northern Ireland) in order to gain insight into how long *C. sylvestre* has been in Ireland. We downloaded records of millipedes between 1970 and 2016 from the Irish National Biodiversity Data Centre (NBDC, http://www.biodiversityireland.ie/ [downloaded 6 October 2017]). Millipede records held by the NBDC primarily came from the BMIG recording scheme (Biological Records Centre, 2017; Lee, 2006), but also included records submitted directly to the NBDC. To evaluate the completeness of millipede recording in Ireland, we estimated species richness using the improved Chao2 estimator (Chui, Wang, Walther & Chao, 2014) and the incidence-based coverage estimator (ICE) (Lee & Chao, 1994), and we graphed species accumulation curves showing the cumulative number of species detected as records were added chronologically and in random order. We estimated species richness and made species accumulation curves for two different time periods (1970 to 1985 and 1986 to 2005), and for both the entire island of Ireland and for the area within 20 km of the new location for *C. sylvestre*. The duration of the two periods during which we estimated species richness was different (14 years for the first period and 20 years for the second period) becauses the two periods were selected based on distinct periods of intensive recording effort. Analyses were conducted in the R statistical programming software, version 3.6 (Chao, Ma, Hsieh, & Chiu, 2016; R Core Team, 2020; Wickham, 2017).

## Results

### New records of Chordeuma sylvestre

One male millipede was identified as *Chordeuma sylvestre* using Blower (1985). The pointed apex of the coxal pillar of the male was not visible *in situ* as described in Blower (1985), but was clearly visible after dissection (Fig 1). The “bristly” flagellum and the pointed apex of the coxal pillar visible on the disected male gonopod (Fig. 1) appeared to match the drawings in Blower (1985) and ruled out *Chordeuma proximum*. The preliminary identification of *C. sylvestre* was confirmed by Steve Gregory and Paul Lee via iRecord, but Steve pointed out that there were other similar chordeumatid species in the Pyrenees that had not been recorded in the UK or Ireland and were therefore not in Blower (1985) and not on the BMIG website. We therefore used DNA sequencing of the COI gene to confirm the species identification. Details of records for both the male and female specimens are in Table 2.

**Table 1.** Observed and estimated species richness, and the total number of records of millipedes in Ireland during two periods of relatively intensive millipede recording. Species richness was estimated using two methods, the improved Chao2 and the incidence based coverage estimator (ICE). Estimates are given as the estimate with the 95% confidence interval in parentheses.

| Time period | Number of records | Number of species recorded | Improved Chao2 estimated species richness | ICE estimated species richness |
| --- | --- | --- | --- | --- |
| 1971 – 1984 | 1416 | 33 | 36.01 (33.82, 43.99) | 40.02 (34.72, 61.56) |
| 1986 – 2005 | 3356 | 40 | 40.25 (40.02, 43.13) | 40.24 (40.02, 43.75) |

**Table 2.** New records of *Chordeuma sylvestre* C. L. Koch 1847 from the campus of University College Dublin, Co. Dublin, Ireland. Latitude and longitude use WGS84. Grid references use the Irish National grid.

| Date | Latitude | Longitude | Grid reference | Sex | iRecord Record ID | GenBank accession number |
| --- | --- | --- | --- | --- | --- | --- |
| 18 January 2020 | 53.3107 | -6.2220 | O 185 304 | Male | 12916137 | XXXXXX |
| 06 March 2020 | 53.3108 | -6.2219 | O 185 304 | Female | n/a | XXXXXX |

**Figure 1.**
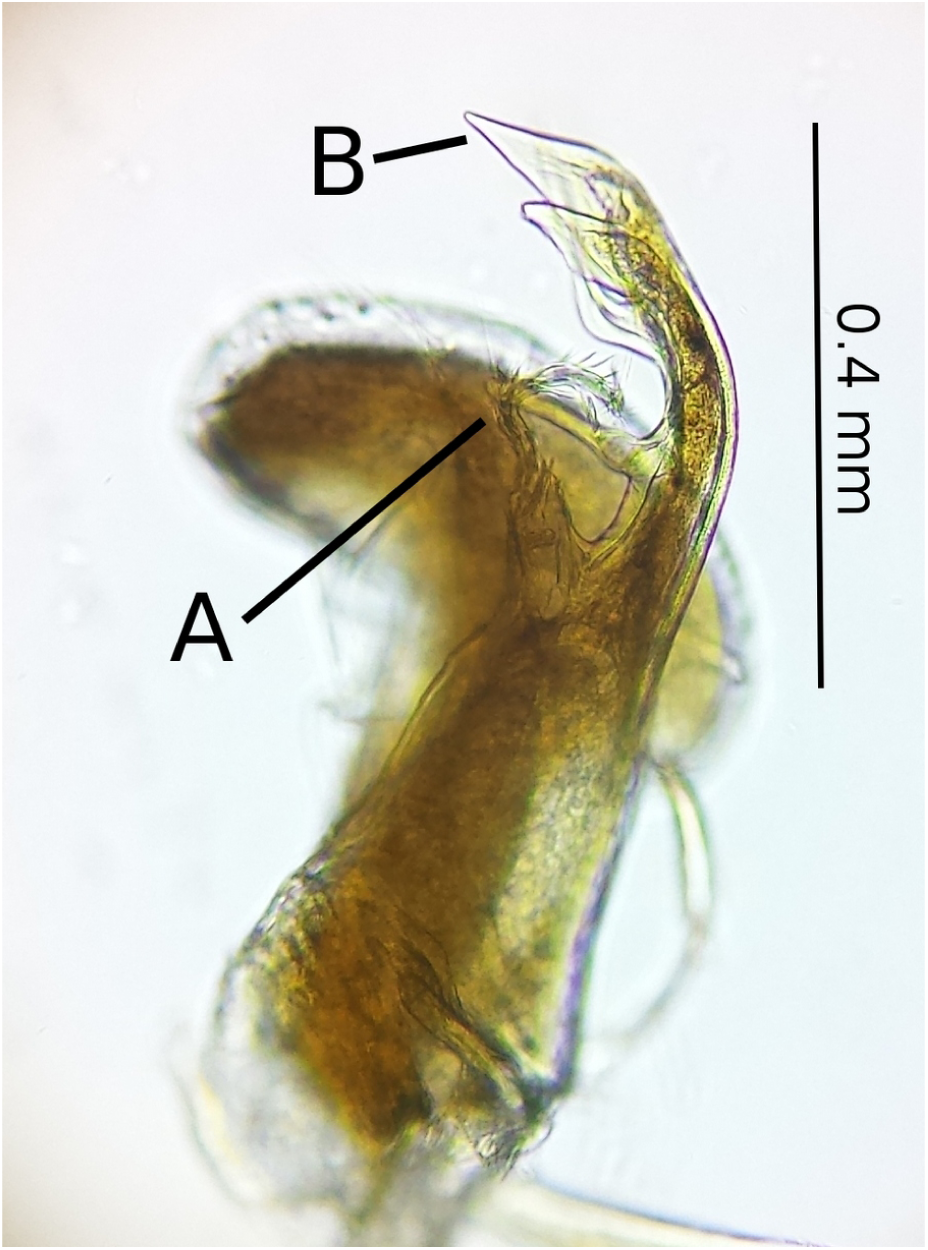
Gonopod of a male *Chordeuma sylvestre* collected in Dublin, Ireland on 18 January 2020, photographed through a microscope. Note the “bristly” flagellum (A) and the pointed apex of the coxal pillar (B).

### DNA barcoding

The two COI sequences (from the male and female specimens) revealed four variable sites in the 662 bp overlap between the two sequences (99.4% match), implying both samples were of the same species, with two different haplotypes present. The BLAST result for the male showed a 99.09% match to a *C. sylvestre* sample collected in Bavaria, Germany (accession number HM888140.1), and the female sequence also showed a 99.09% match to the same sample. Sequences were also checked against the Barcode of Life Database (http://www.boldsystems.org/) (Ratnasingham & Hebert, 2007), which revealed the male sequence to match 99.08% to the same Bavarian sample as in GenBank, in addition to being 99.08% similar to an early release *C. sylvestre* sequence collected in Piedmont, Italy (Fig. 2). The female sample matched 99.39% to the Piedmont sample, as well as 99.08% similar to the Bavaria sample (Fig. 3). Both consensus sequences were uploaded to GenBank (accession numbers: XXXXXX).

**Figure 2.**
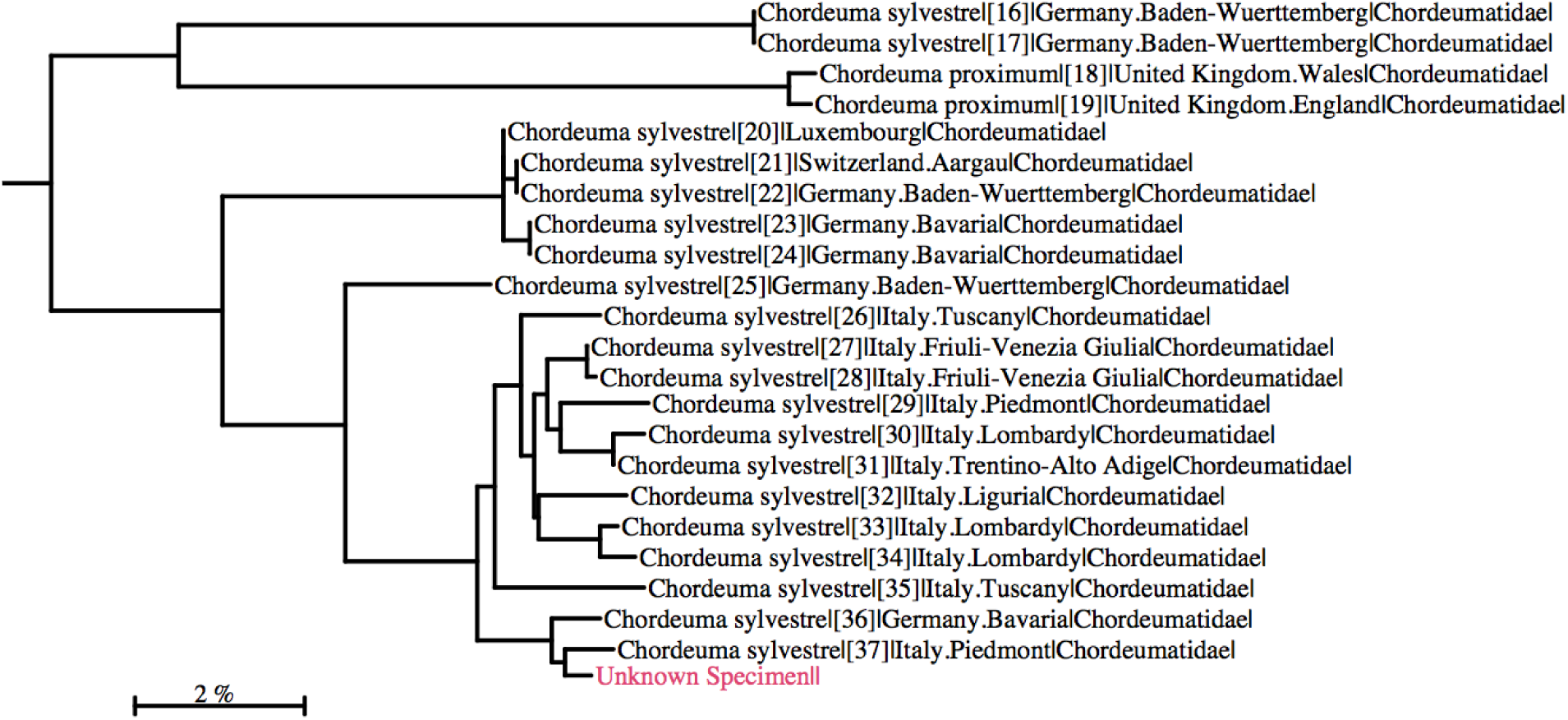
Phylogenetic tree generated for the BOLD results using the Kimura 2 parameter model for the male sample for this study. The sequence generated for the male is highlighted in pink. *Original figure generated by BOLD systems (Ratnasingham & Hebert, 2007) and modified to show only C. sylvestre sequences.*

**Figure 3.**
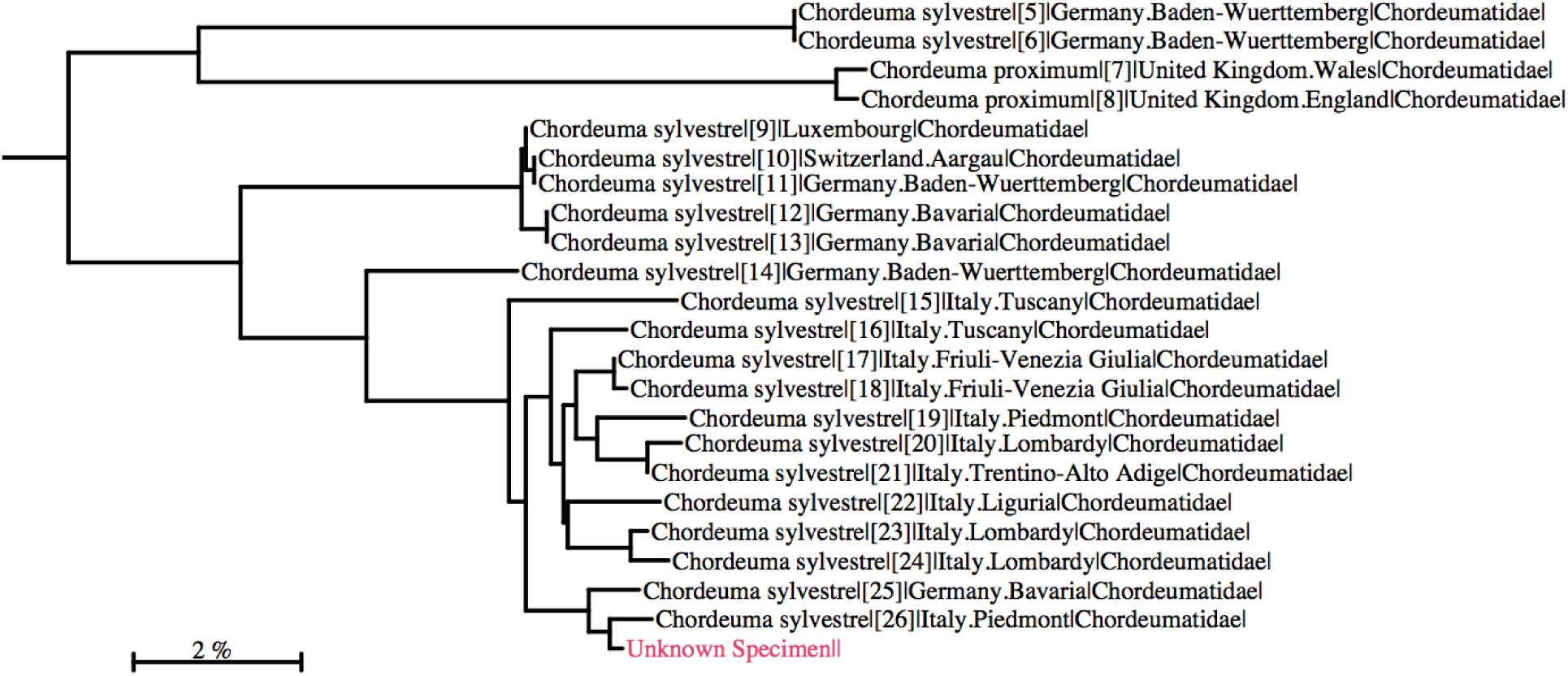
Phylogenetic tree generated for the BOLD results using the Kimura 2 parameter model for the female sample for this study. The sequence generated for the female is highlighted in pink. *Original figure generated by BOLD systems (Ratnasingham & Hebert, 2007) and modified to show only C. sylvestre sequences.*

### Completeness of millipede recording in Ireland

Over 4,800 millipede records have been collected from Ireland since 1970. There is at least one millipede record from 598 hectads (10 km × 10 km grid squares) in Ireland, which is over half of all hectads. However, of the hectads with at least one record, the median number of records per hectad was five, indicating that even hectads with at least some recording remain poorly sampled. Lee (2006) provided a map of the number of records per hectad. The majority of the millipede recording was done in two periods of activity (Fig. 4), from 1971 to 1984 (with a peak in 1978) and from 1986 to 2005 (with a peak in 1994).

**Figure 4.**
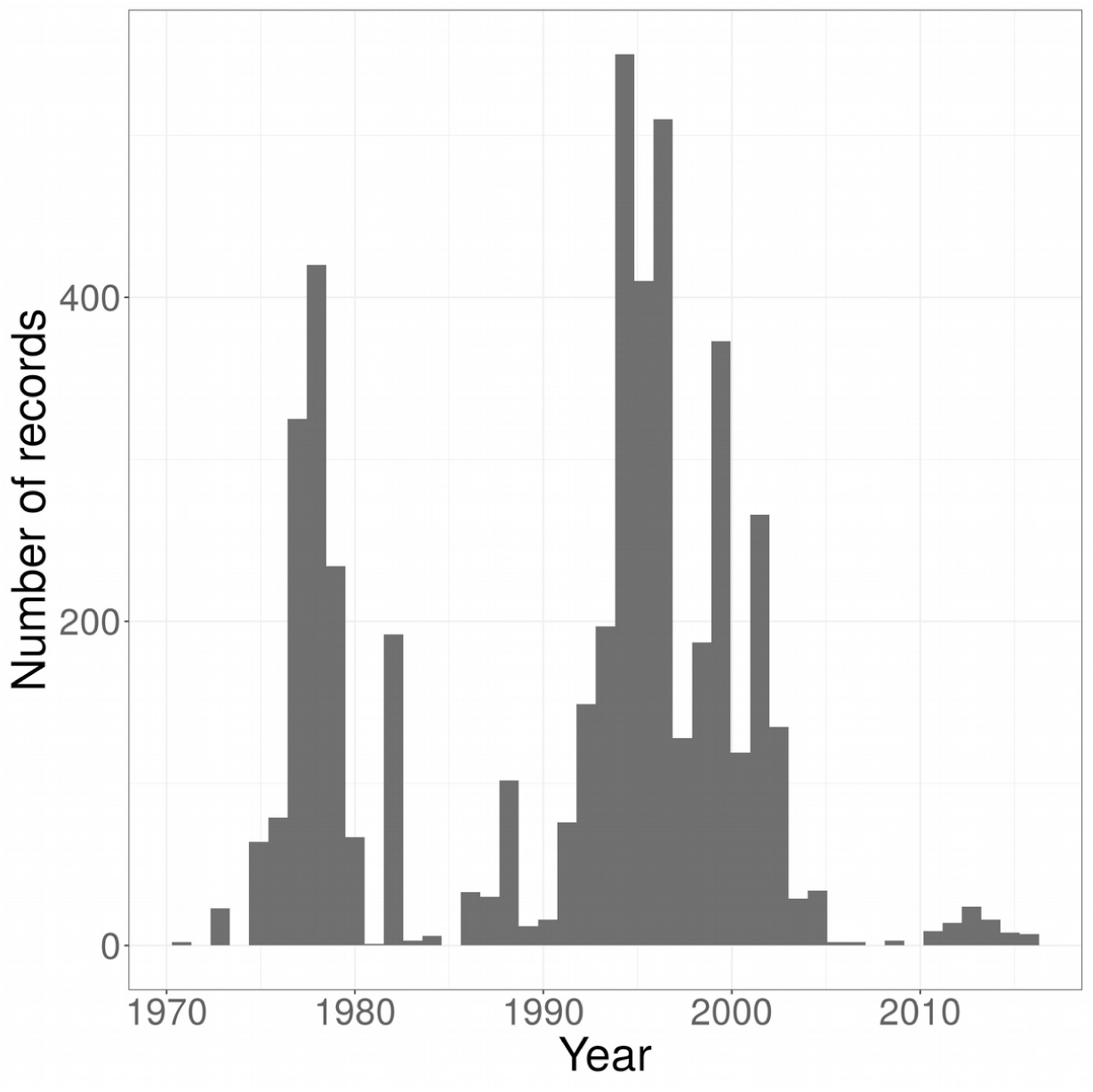
The number of millipede records from Ireland for the years 1970 through 2016. The majority of millipede recording in Ireland occurred during two distinct time periods, from 1971 to 1984 and from 1986 to 2006.

During the period 1971 to 1984, there were 1,416 records of 33 millipede species on the island of Ireland (Tab. 1). Using records from 1971 to 1984, the improved Chao2 species richness estimate was 36.0 (95% CI: 33.8, 43.9) and the ICE species richness estimate was 40.0 (95% CI: 34.7, 61.6) (Tab. 1). During the period 1986 to 2005, there were 3,356 records of 40 species on the island of Ireland. Using records from 1986 to 2005, the improved Chao2 species richness estimate was 40.3 (95% CI: 40.0, 43.1) and the ICE species richness estimate was 40.2 (95% CI: 40.0, 43.7) (Tab. 1). Species accumulation curves showed that the cumulative number of species detected as records were added approached an asymptote in the later period 1986 to 2005 but was still rising steeply during the earlier 1971 to 1984 period (Fig. 5).

**Figure 5.**
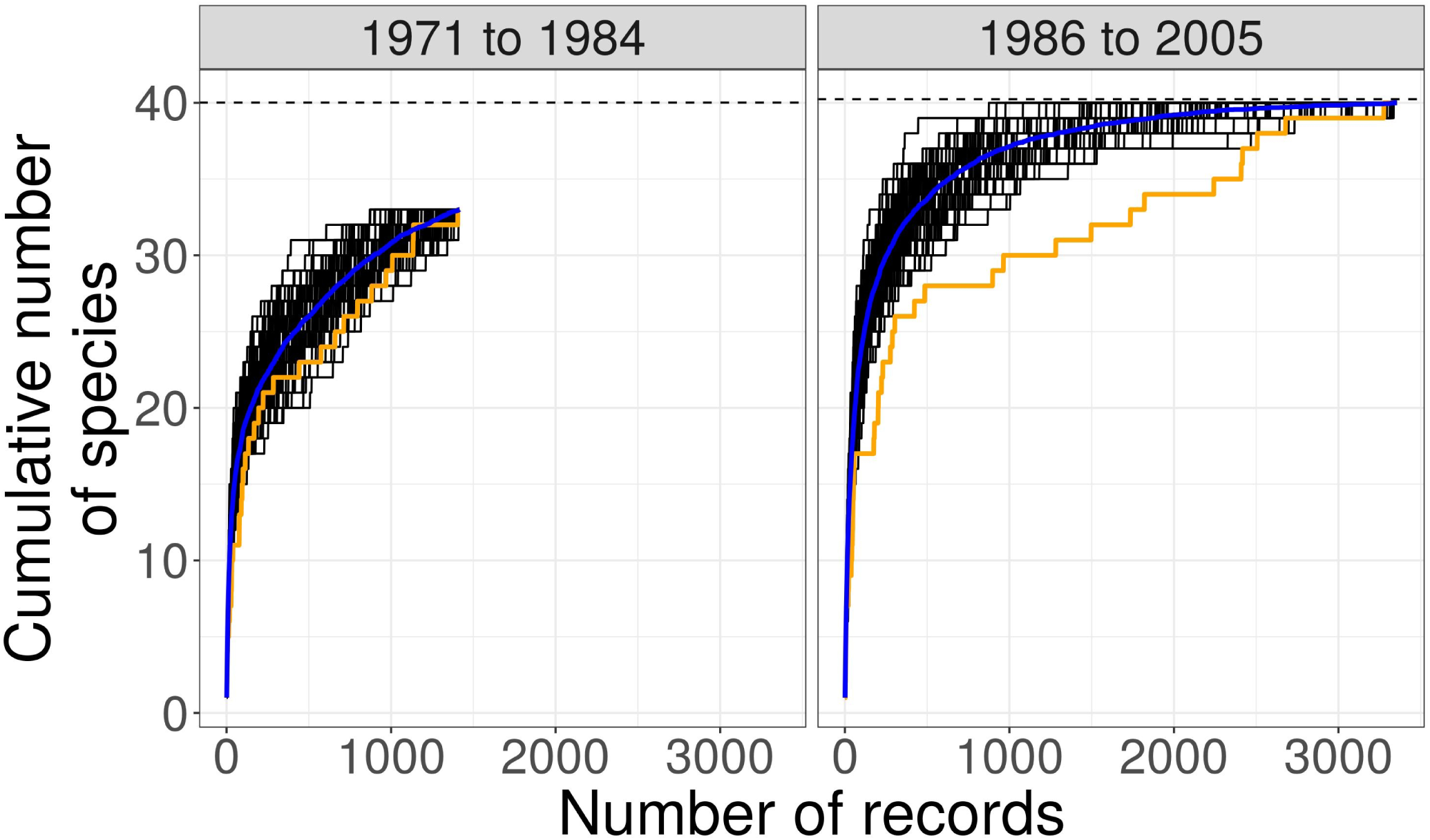
The number of millipede species detected in Ireland as a function of the number of records collected during two periods of intensive millipede recording. The orange line shows the number of species detected as records were added in chronological order. The black lines show number of species detected as records were added randomly for 100 permutations of record order. The blue line shows the mean number of species detected from the 100 randomly permuted orderings of records. The dashed horizontal line shows the ICE-estimated species richness for each time period. During the period from 1971 to 1984, the species accumulation curve did not approach an asymptote and the observed number of species was well below the estimated number, suggesting that there were multiple species present in Ireland but unrecorded during that period. During the period from 1986 to 2005, the species accumulation curve slowed and appeared to approach an asymptote, and the observed number of species was close to the estimated number, suggesting that there were few species present but unrecorded in Ireland during that period.

The species accumulation curves and species richness estimates suggested that the list of species was far from complete from 1971 to 1984, with multiple species present but unrecorded in Ireland. The difference between the observed number of species (33) and the expected species richness (36 species using the improved Chao estimator or 40 using ICE) suggests that there were at least three to seven millipede species present but unrecorded in Ireland during the first period of intensive millipede recording from 1971 to 1984. During the second period of intense recording, from 1986 to 2006, over twice as many records were collected, and, unsurprisingly, the recorded species list seems to have been more complete. The estimated number of species present in Ireland from 1986 to 2006 is the same as the observed number (Tab. 1). The species richness estimates are minimum estimates, so it is possible that there were more than 40 species present during that time period. Nevertheless, the much closer match between the observed and estimated species richness suggests that the species list from 1986 to 2005 was relatively complete.

Within 20 km of the location of the new Dublin *C. sylvestre* records, steeply rising species accumulation curves from both the 1971 to 1984 and 1986 to 2005 time periods indicated that there were many species present but unrecorded in the area during both time periods (Fig. 6).

**Figure 6.**
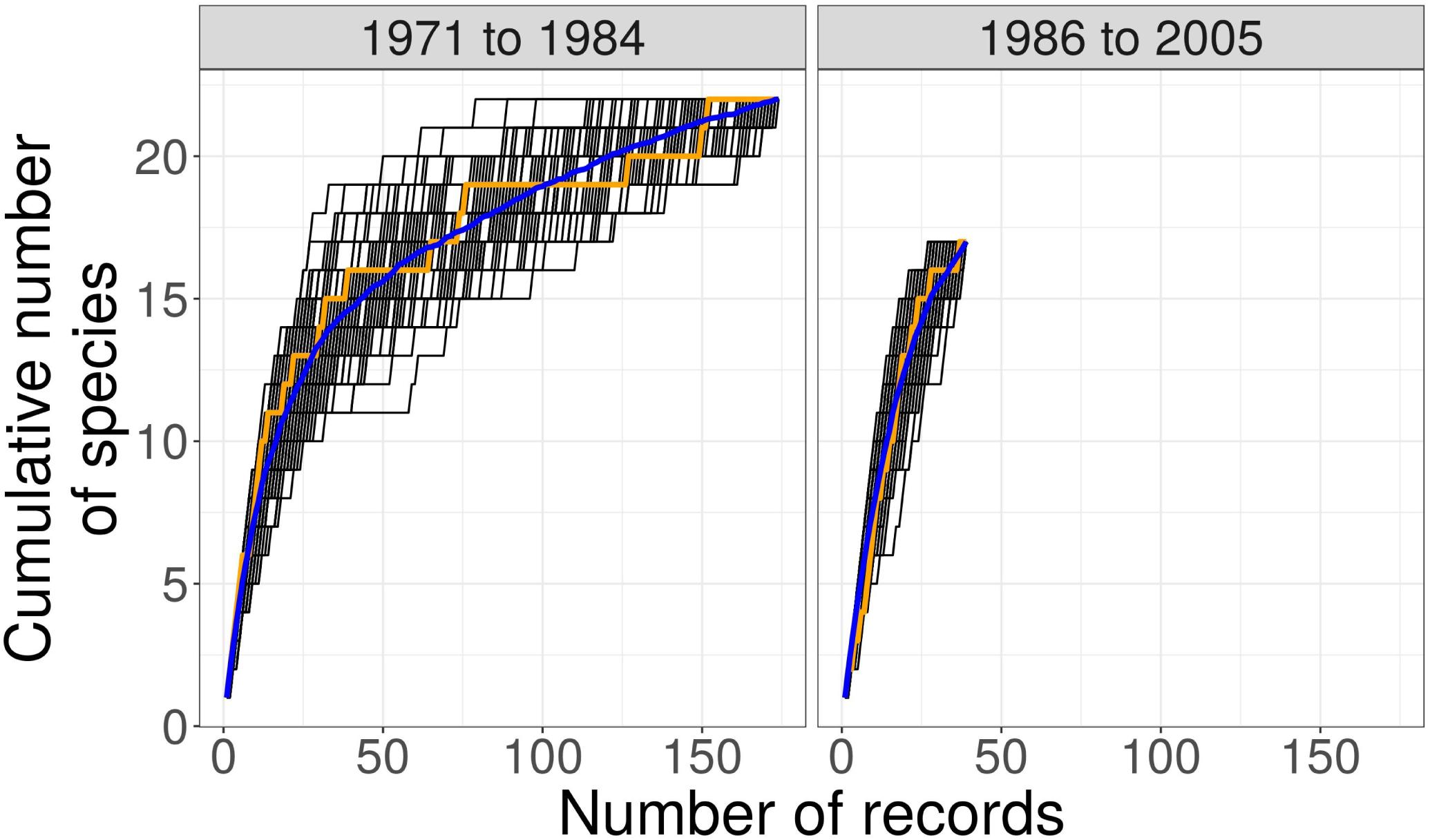
The number of millipede species detected within 20 km of the new *C. sylvestre* location in Dublin, as a function of the number of records collected during two periods of intensive millipede recording. The orange line shows the number of species detected as records were added in chronological order. The black lines show number of species detected as records were added randomly for 100 permutations of record order. The blue line shows the mean number of species detected from the 100 randomly permuted orderings of records. The species accumulation curves did not approach an asymptote in either time period, suggesting that there were many species present but unrecorded in the Dublin area during both time periods.

### Other species recorded from the same site

Other species recently recorded by WG from UCD campus in the same area where the *C. sylvestre* specimens were collected include: *Glomeris marginata* (Villers), *Brachydesmus superus* Latzel, *Polydesmus angustus* Latzel, *Polydesmus coriaceus* Porat, *Ophiodesmus albonanus* (Latzel), *Blaniulus guttulatus* (Fabricius), *Boreoiulus tenuis* (Bigler), *Ophyiulus germanicus* (Verhoeff), *Ophyiulus pilosus* (Newport), *Cylindroiulus punctatus* (Leach), and *Cylindroiulus britannicus* (Verhoeff). This is almost certainly not a complete list and continued recording will likely reveal more species from UCD.

## Discussion

Millipede records from 1986 to 2005 appear to have provided a nearly complete list of millipede species in Ireland during that time. The species richness estimators we used provide only minimum estimates of the number of species present – true number of species could be higher than the estimates. Because recording effort in the Dublin area (within 20 km of the new *C. sylvestre records*) has never been sufficient to compile a nearly-complete species list for that local area (Fig. 6), we cannot rule out the possibility that *C. sylvestre* has been present in Dublin for decades or longer. However, the location of the new records in an urban area, on the campus of a University that gets many international travelers and has relatively extensive landscaping, makes it seems more likely that *C. sylvestre* colonized Ireland within the past few decades. Another millipede species recently added to the Irish list, *Ophyiulus germanicus* (Verhoeff), was found in 2019 in Northern Ireland (Anderson, 2019) and on UCD campus (Gaul, 2020), and was subsequently discovered to have been in samples collected in Northern Ireland as far back as 2015 but mis-identified as *Tachypodoiulus niger* (Leach) (Anderson, 2019). It seems unlikely that *C. sylvestre* has been hiding undiscovered in collected samples in the same way that *O. germanicus* was. The most similar species to *Chordeuma sylvestre* known from Ireland, *Chordeuma proximum* Ribaut, has a relatively limited range in Ireland (Lee, 2006). There is therefore little opportunity for *C. sylvestre* to be mis-identified as a similar-looking, widespread species. On the other hand, morphological identification of *C. sylvestre* (and the similar *C. proximum*) requires an adult male specimen. Blower (1985) reported that only juveniles of *C. sylvestre* were found in August at sites in Great Britain, suggesting that adults may be difficult to find in Ireland during late summer and autumn, though GBIF (2020) has records of *C. sylvestre* from all months of the year in continental Europe. This unavailability of adults at certain times of year could have contributed to *C. sylvestre* remaining undetected in Ireland.

The known locations for *C. sylvestre* in Scotland are from gardens, where Gregory (2016) suggested it was introduced. Lee (2006) and Kime (2001) noted that populations of *C. sylvestre* in Cornwall might be part of the natural range of the species, which is found in northern France. The Dublin and Scottish locations appear even more disjunct from the native range than do the Cornish locations, supporting the idea that the species has been introduced to Dublin. There are no millipede species in Ireland that seem to be strongly limited to Dublin because of environmental conditions, and there is no reason to think that *C. sylvestre* in Ireland would be limited to Dublin by environmental constraints.

With the discovery of *C. sylvestre* in Dublin, at least four additional millipede species have been discovered living in the wild in Ireland since the last period of intense recording ended in 2005, the other three species being *Polydesmus asthenestatus* Pocock (Anderson, 2015), *Cylindroiulus apenninorum* (Brölemann) (Anderson, 2018), and *O. germanicus* (Anderson, 2019). All of these species are somewhat synanthropic in Ireland and it seems likely they have been introduced to Ireland through the movement of plants or soil. At least one other species (the “Ikea millipede” *Xenobolus carnifex*) was found in a potted plant in Dublin but not subsequently recorded living in the wild (Barber, 2015).

Only one species, *Adenomeris gibbosa* Mauriès, was recorded in Ireland between 1971 and 1984 but not recorded from 1986 to 2005. It is possible that the species was lost from Ireland sometime after 1984. However, the records from before 1985 were from the Dublin area, and the species accumulation curves we constructed for the Dublin area (Fig. 6) found that millipede recording in the Dublin area was less complete between 1986 and 2005 than during the earlier 1971 to 1984 period. The 1 km grid squares from which *A. gibbosa* was recorded between 1971 and 1984 were not sampled between 1986 and 2005, and there was only a single record collected between 1986 and 2005 from the same 10 km grid squares in which *A. gibbosa* had previously been recorded. We therefore see no evidence that any millipede species were lost from Ireland between the 1971-1984 and 1986-2005 periods, though our ability to detect a loss is limited given that the species list from the earlier period almost certainly did not include many species that were in fact present.

The number of species now recorded living in the wild in Ireland is 44, which is slightly higher than the upper 95% confidence interval of the species richness estimates for the number of species present between 1986 and 2005 (Tab. 1). However, this is not evidence for an increase in the number of millipede species in Ireland, because, while we can detect apparent additions of species, as reported here, the very incomplete recording since 2005 means that it is impossible for us to detect any loss of species since 2005. A third period of millipede recording, equaling or exceeding the recording effort from the 1986 to 2005 period, would give a better picture of whether the total number of millipede species in Ireland is changing, and would provide information about whether the addition of new species discovered since 2005 has been balanced by the loss of other species.

Identification of millipede species based on genetic evidence overcomes the limitation of adult males not being available during some seasons, as identification is possible for specimens of any age and sex (Savolainen *et al*. 2005; Spelda, Reip, Oliveira-Biener & Melzer, 2011). The cost of identifying species using COI barcoding is relatively low (approximately €9 per specimen if equipment is already available), and the skills and facilities for doing so are within reach even of undergraduate students at many universities.

The presence of two unique mitochondrial haplotypes implies at least two maternal lineages present in Dublin, which rules out colonization from a single pregnant female or a clutch of eggs from one female. We cannot rule out the possibility that *C. sylvestre* was introduced to Ireland from one of the British locations, because there were no sequences available in BOLD from the populations in Great Britain. The COI sequences of the Dublin *C. sylvestre* specimens clustered with a specimen from the Piedmont region in Italy and a specimen from Bavaria in Germany. However, there were few COI sequences available from the northern part of *C. sylvestre’s* range (e.g. no sequences from Belgium or the Netherlands), and no sequences available from France. As more COI sequences from the northern and western parts of the range become available, it may be possible to attribute the source of the Dublin population to the region near the Alps if the Bavarian and Piedmont sequences remain the closest matches.

DNA barcoding of arthropods is becoming an increasingly useful tool for invasive species identification (Armstrong & Ball, 2005; Madden *et al.*, 2019), for narrowing the list of possible source populations (Harris *et al.*, 2017) and – in addition to sequencing of other nulcear loci – for the reconstruction of dispersal and colonization patterns (Guillemaud *et al.*, 2010). One potential drawback is that using genetics to discover a source population requires that the source population has itself been sequenced. This is problematic for poorly recorded taxa such as millipedes that are dispersed anthropogenically, because there will likely always be undiscovered and unsequenced populations. However as projects such as the Barcode of Life Database continue to increase the size of the COI database, this is issue will become less limiting.

## Acknowledgments

We thank Steve Gregory for assistance with identification based on photos of the male specimen, and Paul Lee for permission to use the BRC millipede dataset. Jörg Spelda and Thomas Wesener helpfully answered questions about *C. sylvestre* COI sequences on BOLD. We thank Declan Doogue for informative discussion about collecting techniques and the distribution of millipedes in Ireland. We thank Jon Yearsley and Jens Carlsson for comments on the manuscript and the Area 52 research group at University College Dublin for the use of their lab and materials for DNA extraction. This publication emanated from research supported in part by Science Foundation Ireland (grant number 15/IA/2881), the Marine Institute Cullen fellowship (grant number: CF/17/04/01) and the Crawford Hayes bursary.

